# Sensitivity to Pain Expectations: A Bayesian Model of Individual Differences

**DOI:** 10.1101/176172

**Authors:** R. Hoskin, C. Berzuini, D. Acosta-Kane, W. El-Deredy, H. Guo, D. Talmi

**Affiliations:** Division of Neuroscience and Experimental Psychology, University of Manchester, Manchester, UK; Centre for Biostatistics, University of Manchester, Manchester, UK; Princeton University, Princeton, USA; School of Biomedical Engineering, University of Valparaiso, Chile

**Author notes:** Correspondence: Deborah Talmi, Division of Neuroscience and Experimental Psychology, Faculty of Biology, Medicine and Health, University of Manchester, Oxford Road, Manchester, United Kingdom, M13 9PL. Joint first authors (R. Hoskin, C. Berzuini).

**Keywords:** Pain, Expectation, Uncertainty, Bayes, Perception, Placebo effect

## Abstract

The thoughts and feelings people have about pain (referred to as ‘pain expectations’) are known to alter the perception of pain. However little is known about the cognitive processes that underpin pain expectations, or what drives the differing effect that pain expectations have between individuals. This paper details the testing of a model of pain perception which formalises the response to pain in terms of a Bayesian prior-to-posterior updating process. Using data acquired from a short and deception-free predictive cue task, it was found that this Bayesian model predicted ratings of pain better than other, simpler models. At the group level, the results confirmed two core predictions of predictive coding; that expectation alters perception and that increased uncertainty in the expectation reduces its impact on perception. The addition to the model of parameters relating to trait differences in pain expectation, improved its fit, suggesting that such traits play a significant role in perception beyond those expectations triggered by the pain cue. When model parameters were allowed to vary by participant, the model’s fit improved further. This final model produced a characterisation of each individual’s sensitivity to pain expectations. This model is relevant for the understanding of the cognitive basis of pain expectations and could potentially act as a useful tool for guiding patient stratification and clinical experimentation.

## 1. Introduction

The experience of pain, like all other sensory experience, is a result not only of the objective reality, such as the degree of tissue damage, but also of the sufferer’s beliefs about pain. Conscious and unconscious thoughts and beliefs that people have about imminent pain are referred to as ‘pain expectations’ (Schrooten, Vlaeyen, & Morley, 2012). The effect of pain expectations on pain experience is evident in both laboratory and clinical experiments (e.g. Atlas & Wager, 2012; Bingel et al., 2011; Colloca, & Benedetti, 2006; Peerdeman et al., 2016; Tracey, 2010). In addition, neural correlates of the effect of expectation on pain perception have been established (Brown., et al 2008a; Ploghaus et al., 1999; Seymour et al., 2004; Tracey, 2010; Wager et al., 2004; Watson et al., 2009). Despite these advances there remains a significant gap in our knowledge regarding the specific cognitive processes that underpin pain expectations.

Predictive coding provides the dominant theoretical framework for understanding the effects of expectation on perception (Clark, 2013, Friston, 2003), including pain perception (Buchel, Geuter, Sprenger, & Eippert, 2014; Tabor, Thacker, Moseley, & Körding, 2017; Van den Bergh, Witthöft, Petersen, & Brown, 2017). Predictive coding stipulates that perception is biased towards the expected level of pain, and that this bias will be stronger when the expectation is more certain, since expectation uncertainty causes the suppression of top-down, prior-driven signals, leading to greater importance being placed on the bottom-up sensory input. A number of studies confirm the specific predictions of this theory for pain perception, beyond the established biasing effect that pain expectations exert on pain experience. For example the effect of pain expectations is enhanced when expectations are more precise (Brown, Seymour, Boyle, El-Deredy, & Jones, 2008b; Colloca, Petrovic, Wager, Ingvar, & Benedetti, 2010). Likewise formal Bayesian models provide a good fit to pain reports in placebo analgesia studies (Anchisi, Zanon, Plaghki, Kirsch, & Claggett, 2015; Jung, Lee, Wallraven, & Chae, 2017) and to neural data from the anterior insula (Geuter, Boll, Eippert, & Büchel, 2017).

Although predictive coding guides much of the research on the effect of expectations on perception, some findings appear to contradict its prediction concerning the impact of the precision of the expected pain. Watkinson et al., (2013) delivered two different distributions of pain stimuli, a unimodal and a bimodal distribution. The two distributions had the same mean pain level, but the bimodal distribution had a larger variance. According to predictive coding theories the mean expected level of pain should be the same in the two conditions, but the influence of this expectation on pain experience should be lower in the bimodal condition, as the larger variance of the bimodal distribution should cause greater expectation uncertainty. In direct contradiction with predictive coding, and in support of the alternative, Range Frequency Theory (Parducci., 1963), Watkinson et al., found that the expected mean level of pain had a stronger influence on the rating of three target stimulations in the bimodal condition, despite the greater uncertainty generated by the larger stimulation variance. Likewise, Yoshida et al., (2013) altered participant’s expectations relating to upcoming pain stimulation by presenting fictitious pain ratings of the stimulus prior to their delivery. The mean and variance of the distribution of these ratings were manipulated. They found that increasing pain uncertainty contributed independently to increased pain experience, in contradiction with predictive coding. In addition, it is not clear from Yoshida et al. data whether uncertainty significantly modulated the bias induced by the mean of the fictitious pain ratings, as it should do according to the predictive coding framework. These findings suggest that further work is needed before predictive coding is accepted as a viable framework for understanding pain perception.

Over and above the conflicting findings around the impact of expectation uncertainty, we also know very little about how specific processes that underlie pain expectations are integrated. Unpacking the construct of pain expectations to its underlying information processing mechanisms requires a statistical model of the pain perception process to be formulated which accommodates parameters that describe different facets of pain expectations. Furthermore, in order for such a model to be useful in understanding an individual’s response to both pain and placebo analgesia, it needs to be capable of predicting the effect of pain expectations not just at a group level, but also at an individual level, and not only qualitatively, but also quantitatively. For example such a model would need to be able to identify individuals that experience high levels of pain because they have a trait-like bias to expect high levels of pain (i.e. always expecting high pain, independent of context), and to distinguish such individuals from those who experience high pain because they are highly pessimistic in assessing the information they are given about a treatment, or because they are over-confident in their mildly-pessimistic expectations, so that they rely less on their sensory data. That level of understanding is necessary to allow an individual’s response to treatment to be predicted, thus providing the basis for a tool that can support clinical decision making.

This paper details the construction and testing of a mathematical model of the impact of pain expectations on pain perception using experimental data garnered from a novel predictive cue task. The purpose of this model construction was threefold. Firstly, we wished to assess the effect of expectation uncertainty on pain perception in light of the conflicting past results mentioned above. Our second objective was to identify, using the experimental data, a number of putative cognitive processes that give rise to the umbrella term ‘pain expectations’. Finally, leading on from the second objective, we wished to assess whether a model could be constructed that would enable individuals to be distinguished based on the aforementioned cognitive processes.

We collected two sets of experimental data via two separate experiments using independent samples. A series of increasingly complex statistical models were constructed and their performance was compared using the data from the first experiment. The simplest model (Model 1), where the pain participants experienced was influenced only by the actual, delivered pain, served as a baseline to compare to five other models, which successively included additional facets of pain expectations (Table 1). The multi-modal model (Model 2) represented pain expectations with a multi-modal distribution, which peaked around each of the possible pain levels that participants could expect based on the cues they were given. In the ‘Mean-only model’ (Model 3) pain expectations were assumed to correspond to the mean of the expected pain (i.e. the average of the possible pain levels indicated by the cue). The ‘Mean-and-variance model’ (Model 4) was inspired by predictive coding, and took into consideration the variance of expected pain, a function of the discrepancy between the possible pain levels indicated by the cue. Like Model 3, Model 4 also allowed pain experience to be affected by the mean of the expected pain, but here the impact of mean expected pain was modulated by its uncertainty. Finally the ‘Full model’ (Model 5) additionally included the effect of cue-independent pain expectations in addition to the cue-dependent expectations used in Models 2, 3 and 4. In this formulation, cue-dependent pain expectations relate to those triggered on each trial by the cue, while cue-independent expectations capture more stable, trait-like differences in the propensity to believe that pain will be greater or weaker, independent of the effect of specific local cues. Thus, this model is organised into tiers, such that stable priors contribute to the shaping of more temporary priors triggered by the information provided by the cue. Finally, to satisfy the objective of creating a model that can characterise pain expectations at the individual level, we compared the winning group-level model to an individual-level variant that included individual-level random effects (Model 6). We considered the winning model to be able to usefully characterise pain expectations at an individual level if the ‘individual-level’ variant of the model was significantly better at predicting pain perception than its equivalent group level variant. The second experimental dataset was used for the purposes of conceptual replication; enabling validation of the findings from the first dataset, thus providing evidence that the winning model could predict outcomes in a dataset other than the one from which it was constructed (cf. Maloney & Zhang 2010).

**Table 1:** DIC and LOSO log likelihood values for fixed and variable-parameter models.

|  |  |  | DIC |  | LOSO Log Likelihood |  |
| --- | --- | --- | --- | --- | --- | --- |
|  |  | Model Name | Exp 1 | Exp 2 | Exp 1 | Exp 2 |
| Fixed | 1 | Baseline model | 17007 | 17304 | -8507 | -8657 |
|  | 2 | Multi-modal Model | 16380 | 16793 | -8193 | -8403 |
|  | 3 | Mean-only model | 16178 | 16179 | -8094 | -8096 |
|  | 4 | Mean & variance model | 16138 | 16107 | -8073 | -8063 |
|  | 5 | Full model | 15937 | 15714 | -7981 | -7875 |
| Variable | 6 | Hierarchical model | 15588 | 15047 | -7871 | -7648 |

We hypothesised that the model comparison will provide support for the predictive coding framework. More specifically we predicted, based on neural evidence that the mean of probabilistic cues is computed and utilised in decision-making (Schultz, O’Neill, Tobler, & Kobayashi, 2011), that Model 3 will provide a better fit to the data than Model 2. We also predicted that Model 4 will provide a better fit to the data than Model 3, thus showing that expectation uncertainty significantly affects pain perception; and that the value of model parameter describing the effect of expectation uncertainty will show that increased uncertainty reduces the influence of pain expectations. Based on the vast clinical literature on individual differences in pain catastrophizing (Sullivan, Bishop, & Pivik, 1995) we hypothesised that Model 5 will provide a better fit for the data than Model 4, suggesting that both cue-dependent and cue-independent expectations have separate influences on pain perception. Finally, we hypothesised that when individual-level random effects are added to the winning model this will significantly improve the fit of the model to the data, suggesting that the model is able to characterise sensitivity to pain expectations at an individual level.

## 2. Method

### 2.1 General experimental design

Two experiments were conducted on independent samples. Experiment 2 was conducted for validation, with task delivery modifications introduced to enhance the translational impact of the approach. During each experiment participants performed a ‘pain rating’ task (Figure 1.). In each trial of the task, participants’ expectations regarding the upcoming stimulation were manipulated. This manipulation was achieved by offering the participants a choice between two ‘cues’ expressing different probability distributions for the intensity of the upcoming stimulation in terms of the possible intensities and corresponding probabilities. After selecting one of the cues the participant then experienced a pain stimulation level generated in accordance with the probabilities represented by the selected cue. Finally, the participant was required to rate the intensity of the stimulation they received. This choice mechanism was used (in contrast to just presenting participants with one cue) to ensure that the participants both attended to, and understood, the pain probability cues they were presented with. The choice mechanism also acted to give the participants a sense of control over the upcoming pain, so as to minimise the possible influence of learned helplessness on their pain experience (Bhat et al., 2010). The trials were designed in such a way that one cue (herein referred to as the ‘target’) was always preferable to the other (herein referred to as the ‘lure’) in terms of expected pain intensity. Trials where the participants chose the lure (or failed to make a choice at all) were discarded from the data analysis as in such instances it was not certain that the participants had understood the cues. The targets were created in accordance with a design that allowed a systematic manipulation of the relevant aspects of pain expectation.

**Figure 1.**
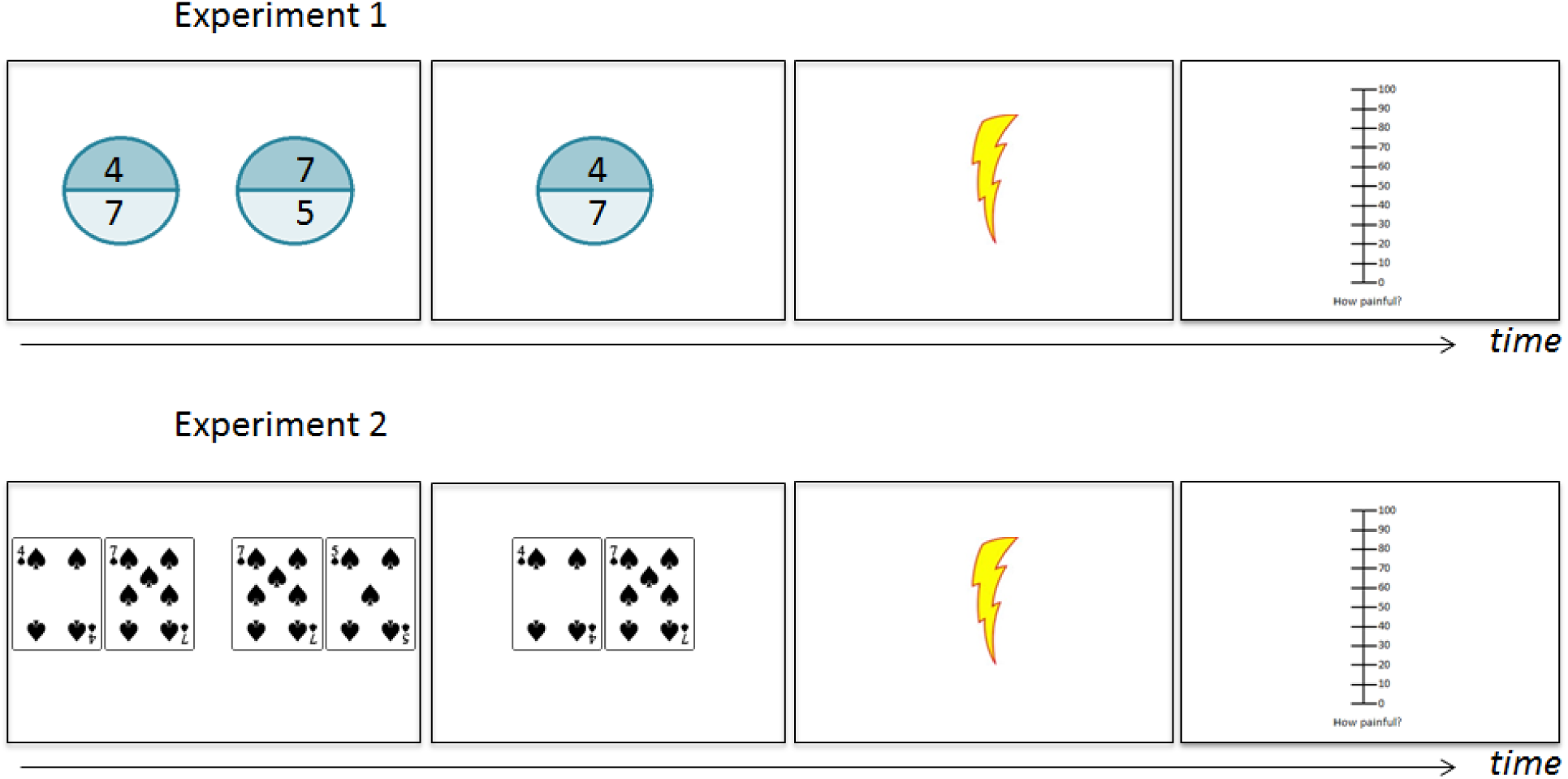
Schematic of one experimental trial in experiments 1 (top) and 2 (bottom). Each trial began with a cue selection phase (left panel) where the participant had to choose between two cues. Participants who understand these cues should choose the left, target cue to minimise the chances of receiving strong pain. Once a cue was selected it appeared on its own for 2s (second panel). This selected cue was assumed to influence participants’ pain expectations. Here the selected cue gives the participant 50% chance of receiving pain level ‘4’ and 50% of receiving pain level ‘7’. With this cue selected the expected pain level for a participant who is not particularly anxious or overly optimistic is 5.5. Participants then received a painful electric stimulation corresponding to one of the levels depicted on the selected cue, e.g. a ‘4’ (third panel). Finally the participant rated their pain experience (fourth panel). In Experiment 1 the size of the portion on which the number was displayed corresponded to the probability of receiving pain of that level. In Experiment 2 all probabilities were fixed to 50%.

The task utilised here is novel in the context of pain research. However, it has been used extensively to examine how people value financial outcomes (Talmi, Atkinson, & El-Deredy, 2013) as well as taste and pain outcomes (Hird et al., 2017). It differs from placebo paradigms because the manipulated quantity is the individual’s expectation about the impending pain, rather than about the efficacy of treatment. One particular feature of this task (in contrast with placebo and other cue-based paradigms) is the avoidance of deception. In placebo paradigms participants are told that an ineffective intervention (e.g. an inert cream) is in fact a proven treatment (e.g. an active analgesic). In current cue-based paradigms (e.g. Atlas et al., 2010), cues intentionally mislead participants, for example informing them that an impending pain stimulus will be high intensity, when it may in fact be of moderate intensity. We avoided deception firstly because the integrity of data resulting from deception paradigms is reliant on the participant remaining naïve to the deception throughout the experiment, and secondly because deception is problematic as far as translation into clinical settings is concerned. Files needed to run the experimental paradigms, the analysis code and the complete datasets are available to download from BioRxiv.

### 2.3 Experiment 1: Method

#### 2.3.1 Experimental design

The top panel of Figure 1 shows an example of a trial from Experiment 1. In this example the ‘target’ cue presented a 50% chance of receiving level 4 pain and 50% chance of receiving level 7 pain. This target therefore provided an expected pain intensity of 5.5, and is therefore preferable to the ‘lure’ cue, which presents a 50% chance of receiving level 5 pain and 50% chance of receiving level 7 pain (expected intensity = 6). The cues were designed to provide a balanced manipulation of the following variables across the sequence of trials experienced by each individual: the expected value of the pain (the mean: 4.5, 5.5), the number of different pain levels that could potentially be delivered (2, 4), the prediction error (the difference between expected level of pain and intensity of the delivered stimulation: 0.5,1.5) and the sign of the prediction error (positive or negative). Using the trial depicted in Figure 1 as an example, if the target cue was selected and the level 7 pain subsequently delivered, then the prediction error would be +1.5 (7 - 5.5). The positive sign of the prediction error indicates that the pain level that was experienced was higher than the average pain level participants would expect on the basis of the cue they selected. In summary, the experimental corresponded to a 2 (mean expected pain: 4.5, 5.5) × 2 (number of cue options: 2, 4) × 2 (direction of the prediction error: positive, negative) × 2 (the size of the prediction error: 0.5, 1.5) design.

#### 2.3.2 Participants

Sixteen undergraduates from the Manchester University School of Psychological Sciences (14 female, Mean age 19.6, σ = 1.36) participated in the study for course credit. Participants were excluded if they had a history of psychiatric or neurological disorders. All participants achieved the criterion performance level of 85% correct responses (defined as choosing the target cue during the pain rating task). The study received ethical approval from the North West 6 Research Ethics committee (Greater Manchester South).

#### 2.3.3 Materials

Acute experimental pain was delivered to the participants using electric stimulation, which has been shown to produce effects of expectations that are equivalent to those produced with the more typical laser stimulation (Hird, Jones, Talmi, & El-Deredy, 2018). Compared to laser pain, which is used more often in the laboratory, the electric stimulation method is safe to use repeatedly and easier to implement in clinical settings. The electrical stimulations were delivered to the back of the right hand via a ring electrode built in-house (Medical Physics, Salford Royal Hospital) attached to a Digitimer DS5 Isolated Bipolar Constant Current Stimulator (www.digitimer.com). To ensure adequate conductance between the electrode and the skin, the back of each participant’s hand was prepared with Nuprep Skin Preparation Gel and Ten20 Conductive Paste prior to the electrode being attached. The experimental paradigm itself was delivered via a laptop using Cogent2000 on a Matlab platform (www.Mathworks.com). The inputs to the DS5 machine were sent from Matlab via the Spike2 software (www.ced.co.uk) and a 1401plus data acquisition interface (www.ced.co.uk).

Cues consisted of charts which were 240 pixels in diameter and which used colours that were equiluminant (thus making each chart equiluminant). These charts used combinations of the 5 pain levels (3,4,5,6,7) and 6 different proportions (25%, 33%, 50%, 67%, 75%, 100%) to manipulate aspects of the pain expected (Supplementary Table 2.)

#### 2.3.4 Procedure

Participants were given an information sheet prior to the study informing them of the justification for the study and of the use of electrical stimulation. On arriving at the laboratory participants were first asked to sign a consent form, and then to confirm that they had read the information sheet. The electrode was then attached to the back of the participant’s right hand. Once the electrode was attached the participants undertook a pain calibration procedure. This procedure was necessary firstly to ensure that the participant could tolerate the stimulations, and secondly to ensure that the stimulations were psychologically equivalent across participants. Once the pain calibration procedure was complete participants were given instructions relating to the pain rating task. Participants were then given 6 practice trials of the pain rating task before undertaking the task proper.

##### 2.3.4.1 Pain calibration

During this procedure participants received a series of stimulations, starting from 0.2V, and incrementing by 0.2V at each step. Participants rated each stimulation on a scale from 0 – 10 where a score of 0 reflected not being able to feel the stimulation, 3 reflected a stimulation level that was on the threshold of being painful, 7 related to a stimulation that was deemed ‘painful but still tolerable’ and 10 related to ‘unbearable pain’. The scaling procedure was terminated once the participant reported the level of pain as being equivalent to ‘7’ on the scale. This calibration procedure was performed twice to allow for initial habituation/sensitisation to the stimulation. The voltage levels rated as ‘3’, ‘4’, ‘5’, ‘6’, and ‘7’ on the second run of the calibration procedure were used as the different pain levels to be delivered during the pain rating task. The average voltage used for each pain level are shown in Supplementary Table 1.

##### 2.3.4.2 Pain rating task

Each trial began with a 250ms fixation cross, before two cues, presented as two pie charts, appeared side-by-side on the screen (+/- 160 pixels from the centre: Figure 1). Participants were required to choose between these cues using the mouse. Numbers on the ‘slices’ of each pie chart indicated the level of stimulation that the slice related to, while the size of the slice depicted the probability of the stimulation level being delivered if the cue was selected. Participants had up to 8 seconds to make their selection. The target cue was always clearly preferable to the lure in terms of expected pain value (see General Methods). If the participant failed to make any selection within the 8s time limit then the lure cue was automatically selected. The selected cue was then presented alone in the centre of the screen for 2 seconds, at which point the cue disappeared and one of the pain levels depicted on the cue was delivered to the back of the hand according to the probabilities depicted on the cue. Finally a visual-analogue scale (Hjermstad et al., 2011) ranging from 0–100, appeared 500ms after the offset of the stimulation. Participants were required to rate their experience of the stimulation on this scale using the mouse, with the trial only proceeding once a rating had been given. Participants were told that even though the stimulation they would receive would always be one of those predicted by the selected cue, they should rate the pain they actually felt. The use of a 0–100 rating scale gave the participants sufficient scope to report trial-by-trial differences in pain experience, while preventing them from simply reporting which intensity they thought had been delivered (which would be a possibility if participants were asked to rate their experience using the same 0–10 scale used during the pain calibration task). The starting position of the cursor (on the rating scale) was randomised on a trial-by-trial basis, eliminating systematic biases of the expectations on the motor response. After the rating was provided an inter-trial interval occurred of a length selected randomly from 1000, 1500, 2000, 2500 & 3000ms (average 2000ms).

##### 2.3.4.3 Experimental task

Participants undertook 5 blocks of trials, with 32 trials in each block, giving a total of 160 trials. Each block consisted of a randomised presentation of 24 types of trials, depicted in Supplementary Table 2. In 16 of these the target cue corresponded to one of the 16 experimental conditions according to the 2×2×2×2 experimental design described above. Four additional trial types were required to complete the design (see Supplementary Table 2). Four further trials types, involving target cues which depicted the delivery of pain levels 3, 4, 5 and 6 with 100% probability, were included for the purpose of a manipulation check. A self-timed break was given every 20 trials.

### 2.4 Experiment 2: Method

The design of Experiment 2 was similar to that of Experiment 1. The following changes were made with the potential clinical application of the paradigm in mind. Firstly, sets of playing cards were used as cues instead of pie charts (Figure 1). This allowed the results from Experiment 1 to be (potentially) replicated using cues that may be more readily understood by a clinical audience. Secondly only trial types involving 2 different intensities with equal (50%) probabilities were included in the design. This meant that only 3 aspects of pain expectation were manipulated within participants; the expected pain level (4.5, 5, 5.5), the size of the prediction error (0.5, 1.5, 2), and the direction of the prediction error (positive or negative). This change allowed a reduction to the number of trials appearing in the experiment, and therefore a reduction in the length of the experiment. This was thought beneficial due to the limited time available in the clinic for diagnostic procedures.

#### 2.4.1 Participants

Thirty-two undergraduates from the Manchester University Psychology Department (26 female, Mean age 19.3, σ = 1) participated in the study for course credit. Participants were excluded if they had a history of psychiatric or neurological disorders. The study received ethical approval from the North West 6 Research Ethics committee (Greater Manchester South). Data from three participants was excluded because they did not achieve the criterion performance level of 85%, leaving a sample of 29 (24 female, mean age 19.3, σ = 1).

#### 2.4.2 Materials and procedure

The materials were the same as in Experiment 1, but instead of charts, the cues consisted of spade suit playing cards. Each cue took the form of a pair of playing cards (100×130 pixels in size), with each card corresponding to one slice of the charts used in Experiment 1. During the selection phase the two cards from each pair of cards were presented and pixels from the centre of the screen, with one pair being presented to the left of the screen and the other to the right. After selection, the two cards comprising the selected cue were presented +/- 75 pixels from the centre of the screen.

Participants undertook 5 blocks of trials. Each block consisted of a randomised presentation of 16 types of trials depicted in Supplementary Table 3. In 8 of these, the target cue corresponded to one of the 8 experimental conditions in a 2 (the expected pain level: 4.5, 5.5) × 2 (the size of the prediction error: 0.5, 1.5) × 2 (the direction of the prediction error: positive or negative) experimental design. In 4 trial types the target cue corresponded to one of the 4 experimental conditions in a 2 (the size of the prediction error: 0.5, 1.5) by 2 (the direction of the prediction error: positive or negative) experimental design. We employed these two designs together in order to compare their efficacy, and decide which design was best for future experiments. Four additional trial types involving target cues which depicted the delivery of pain levels 3, 4, 5 and 6 with 100% probability were again included for the purpose of a manipulation check. In total, therefore, each participant undertook 80 trials, with self-timed breaks given every 20 trials. The average voltage used for each pain level during the experiment are shown in Supplementary Table 1.

### 2.5 Statistical analysis

#### 2.5.1 Modelling

The predictive coding framework suggests that our perceptual systems are organised in a hierarchical manner. Within this framework perceptual systems at a high level in the hierarchy hold prior distributions relating to sensory input which are shaped by past experience. Perceptual expectations at any specific point in time arise from these prior distributions. For example, during pain perception, the prior distributions (priors) held at high levels of the hierarchy, are compared to incoming pain at lower levels. Discrepancies between the priors and sensory input are then projected back to the higher level, causing the prior to be updated into a posterior distribution (from which the resulting pain experience arises) while also altering the basis on which subsequent priors will be calculated (Buchel et al., 2014). The models presented in this paper are influenced by this framework, formalising response to pain in terms of a Bayesian prior-to-posterior updating process. Apart from the baseline model, we consider 5 Bayesian models, which are all based on a multiplicative decomposition of the joint probability that conforms to the Bayes theorem, under the stipulated assumptions. These 5 models can be considered Bayesian because they allow a prior distribution representing pain expectation to be updated, on the basis of sensory information, into a posterior distribution representing the final perceptual experience. The models are Bayesian also in that they regard model parameters as random variables. From a statistical standpoint the full version of these Bayesian models is ‘hierarchical’ in that it involves a multilevel Bayesian structure consisting of two levels (tiers) of the model specification:

1. a bottom level where distribution is specified conditional on unknown model parameters
2. a top level which specifies a distribution of the parameters which (in the final model) are allowed to vary from one individual to the next according to an exchangeable population distribution.

For participant *i* during trial *j* we represent the pain stimulation that was delivered as *X_ij_* and the pain they reported as *R_ij_*, both expressed on a 0–100 scale. The ratings were linearly transformed (*R_ij_* → *R*′*_ij_* = *a_i_* + *b_i_* × *R_ij_*) using the pain responses on the 100% probability trials to standardize the responses across participants so as to avoid issues of differential scale use. Thus *a_i_* and *b_i_* were selected to minimize the squared error between participants’ ratings and the delivered intensity. Before the pain stimulation was delivered, participant *i* at the *j*th trial also saw a cue. Let the cue consist of information *Z_ij_*, involving the possible magnitudes 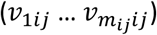 of the incoming pain stimulus, also expressed on a 0–100 scale, and their corresponding probabilities of occurrence, 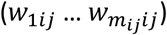. Thus conditional on *Z_ij_*, the intensity of the incoming pain stimulus has mean *q_ij_* and standard deviation *sd_ij_* respectively given by

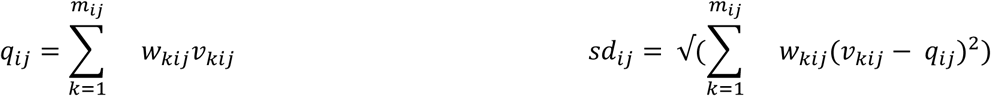

On the *j*th trial, an individual *i* exposed to cue *Z_ij_* represents the predicted intensity of the upcoming pain as a probability distribution P(*R_ij_* | *Z_ij_*). In the terminology of predictive coding, this is the ‘prior distribution’ of the intensity of the upcoming pain. We formulate *P*(*R_ij_* | *Z_ij_*) as a product of two normal distributions, one incorporating cue-independent information and the other incorporating the cue information (see appendix for derivation):

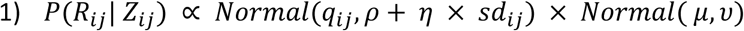
where the symbol ∝ stands for “proportional to” and *Normal(x, y)* stands for the normal distribution with mean *x* and standard deviation y, occasionally truncated to ensure that random variables satisfy their range constraints, whenever appropriate.

The cue-independent *Normal*(*μ*,*υ*) represents a stable, trait-like bias to expect pain of a certain intensity independently of the cue. Higher values of parameter *μ* indicate higher values for the mean of the cue-independent prior (and thus reflect the expectation of higher levels of pain, independent of any cue information). Higher values of parameter *υ* (the standard deviation of the cue-independent prior) reflect greater uncertainty (and therefore lesser influence) of this cue-independent prior. Participants with high *μ* and low *υ* can be thought of as pessimistic about pain. The cue-dependent *Normal*(*q_ij_*, *ρ*+ *η* × *sd_ij_*) represents the effect of the cue information *Z_ij_* on the predicted (pre-stimulus) rating. The parameter *ρ* controls the precision of this prediction. Parameter *η* modulates this precision according to the cue’s variance, such that increasing values of *η* indicate a greater detrimental effect of variance within the cue on the precision of the prediction. The influence of the cue thus increases with lower values of *ρ* and *η*. Together, these parameters determine the extent to which the cue information modulates the prior distribution, and thus the subsequent experience of pain.

The stimulation causes the prior distribution to be updated into a posterior. The posterior distribution P(*R_ij_* | *X_ij_*, *Z_ij_*) governs the pain intensity that the individual experiences and rates. Following from Bayes’ theorem, the posterior is given, up to a proportionality constant, by the product of the likelihood of the stimulation and the prior:

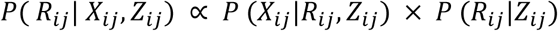

In the appendix we justify the following:

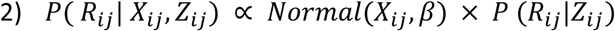

In this term, each predicted value of *R_ij_* is compared with the external sensory information, associating each such value with a measure of ‘surprise’, which our model takes to increase with an increasing distance between *R_ij_* and the actual intensity of the stimulation, *X_ij_*. The value of *ϐ* determines the extent that the perceptual mechanism will tolerate a given disparity between the predicted experience and the actual sensation, with smaller values of *ϐ* implying greater sharpness of the likelihood function around *X_ij_* and hence a reduced tolerance. Plugging Equation 1 into Equation 2 we get the full model:

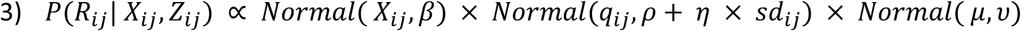

Here the prior is combined with the surprise in accord with the Bayesian formalisation of the integration of multiple sources of evidence. The multimodal prior model (Model specification section below) elaborates the second term of Equation 3 into a mixture of normals

#### 2.5.2 Model specification

Data from each experiment were analysed separately with the following models:

*1. Baseline model*: This model assumes that pain experience depends solely on the pain stimulation. No effect is attributed to pain expectations, whether arising from the cue, or from cue-independent effects of the prior.

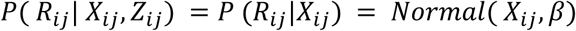

*2. Multimodal prior model*: In this model pain experience depends not only on the pain stimulation, but also on the information in the cue, where that information is represented by a multi-modal weighted mixture of normals. The spread of each normal component of this mixture is modulated by the parameter *ρ*. The normals are centered at each possible magnitude 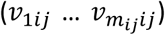 of the incoming pain stimulus and are weighted by their corresponding probabilities of occurrence, 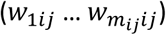.

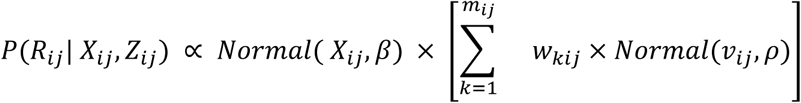

*3. Mean-only model*: As with Model 2, in this model pain experience again depends not only on the pain stimulation, but also on the information in the cue. Unlike Model 2 however, the cue information is represented by the mean of the cue, *q_ij_*, modulated by the parameter *ρ*.

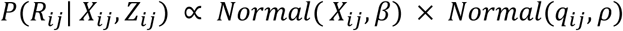

*4. Mean and Variance model*: Pain experience depends not only on the pain stimulation, but also on the mean and variance of the cue. These influence the uncertainty of the prior distribution of *R_ij_* such that cues with higher *sd_ij_*(modulated by *η*) influence the posterior less than precise cues.

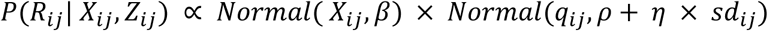

5. *Full model:* In this model pain experience is determined jointly by the pain stimulation and two tiers of expectations: stable cue-independent effects akin to optimism/pessimism trait and effects informed by the cue. Cue-independent effects are represented by a probability distribution with mean *μ* and variance *υ*, which is integrated with the information provided by the cue to form the final prior distribution.

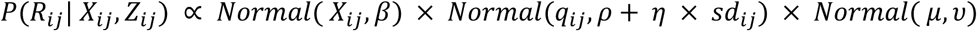

6. *Hierarchical model:* In this model pain experience is also determined jointly by the pain stimulation and two tiers of expectations (as in Model 5), however in this model the parameters are now allowed to vary by participant.

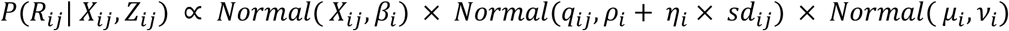

#### 2.5.3 Inference

All the models in our analysis included a specification of the likelihood of the modeled observed quantities *X* and R, given the model unknown parameters and the non-modeled observables, Z, plus a specification of a (possibly uniform) prior distribution for the unknown parameters. Data from each experiment were analysed separately. After each fitting, the goodness of fit of the model to the data was assessed via the Deviance Information Criterion (DIC, Spiegelhalter et al., 2002) which penalises models for increasing complexity. This criterion is sensitive to non-normalities of the posterior distributions of the parameter. An exploration of these posteriors did not reveal departures from normality that might raise concern. At any rate, in order to protect from any anomalous behaviour of DIC, we ranked the models also in terms of a cross-validated measure of performance, specifically, leave-one-subject-out (LOSO) cross-validation (Xu & Huang., 2012). In LOSO, the data for each experiment is partitioned by participant. For each partition the models are fit with the remaining data from the same experiment and a log likelihood of the hold out data is calculated. The DIC-based ranking and the cross-validated ranking were identical.

Using data from Experiment 1, we first compared fixed-parameter models 1–5 where all participants shared the same values for the parameters. In these comparisons the baseline model (Model 1) serves as the null hypothesis, as it assumes that the cue exerts no effect on pain experience, namely, no effect on the prior and thus no effect of expectations of any kind. The comparison between the mean-only model (Model 3) and the multimodal model (Model 2) serves to assess the relevance of the expected value of the cue. Following these comparisons, the same data were analysed with an elaboration of the winning fixed-parameters model, now allowing the parameters *ϐ, ρ, η, µ, and ν*, and to vary across individuals (the ‘Hierarchical’, Model 6). We took the values of these parameters in individual *i* (respectively denoted by *ϐ_i_ ρ_i_*, *η_i,_ µ_i_*, and *ν_i_*,) to be independently drawn from corresponding population-level normal distributions, with hyperparameters independently drawn from flat hyperprior distributions, resulting in a multilevel model. This model allows us to describe the variability of the individual-specific posterior estimates for parameters, and therefore to infer how the parameters vary across the population. Finally, we fitted these models to data from Experiment 2, to provide validation that the findings from the first dataset would generalise to an independent dataset.

As regards parameter estimation, the models, when combined with data, gives rise to a Bayesian posterior distribution for the unknown model parameters. We sampled this posterior by using the Hamiltonian dynamics Markov chain Monte Carlo (HDMCMC) techniques (Metropolis, Rosenbluth, Rosenbluth, Teller, & Teller, 1953; Neal, 2012) incorporated in the program Stan (Stan Development Team, 2014a, 2014b). Any summary of the marginal posterior for any parameters of inferential interest can be reconstructed on the basis of the sampled value for that parameter, generated by the Markov chain after a suitable number of “warm-up” iterations. Initial values for the Markov chains were obtained via variational inference techniques (Wainwright, & Jordan, 2008) based on a parametric approximation of the model posterior.

## 3. Results

The average performance during the cue selection task (i.e. proportion of targets selected) was 96% in Experiment 1 and 97% in Experiment 2, suggesting that participants understood the task well. In both experiments the mean ratings given during the trials where the cue provided 100% chance of a particular stimulation intensity increased linearly as a function of increased stimulation levels, confirming that participants were able to distinguish between the stimulation levels (Supplementary Table 4).

Table 1 summarizes the DIC and LOSO log likelihood values for the fixed-parameter models. Lower DIC values indicate higher likelihood of the data being generated by the model and thus better fit. Higher LOSO values (lower in absolute value) also indicate a better fit. A difference greater than 10 between the DICs for two models is considered strong evidence against the model with the higher score. For Experiment 1, Model 1 fitted the data less well than the other models, which have all assumed that the cue has some effect on the rating. The ‘Mean-only model’ (Model 3) fitted the data better than the ‘Multimodal model’ (Model 2) justifying the use of the mean of the cue as a useful property of the cue information. The best-fitting fixed-parameter model was the ‘Full model’ (Model 5). This finding supports the hypothesis that both cue-dependent and cue-independent biases influence pain perception. When the same models were tested using the data from Experiment 2, the results of the model comparisons were the same (Table 1).

Table 2 summarizes the parameter estimates achieved from fitting Model 5, the winning ‘Full model’. Within this perspective, the finding that the average value of ρ is of smaller magnitude than the average value of β suggests that that the prior distribution for *R_ij_* is biased towards the mean of the cue. These parameters are inversely proportional to the strength of the cue and stimulus information when they are combined to form the posterior in this model. Thus, the parameter values suggest that an increase in the mean of the cue will, on average, and all other variables remaining equal, correspond to an increase in the mean of the rating *R_ij_*. The positive average value of η supports the hypothesis that an increase in the cue variance will, on average, correspond to an increase in the variance of the prior distribution for *R_ij_* and thus a smaller effect of the cue on the perceived intensity, in accordance with predictive coding.

**Table 2:** Estimates of the model parameters arising from the full model. Parameter values from the ‘Full model’ (Model 5), where the parameters are assumed to be identical across the participants. The values in the table represent the mean and standard error (the standard deviation of the marginal posterior distribution) of each estimated parameter, respectively. The corresponding credible interval, defined for each parameter by the mean plus/minus 1.96 times the standard deviation, can be interpreted to contain the true value of the parameter with probability 0.95. The model parameters can be described as follows. ϐ: This parameter controls how likely is a value of R_ij_ conditional on X_ij_, as a function of the distance between the values of R_ij_ and X_ij_. ρ: The increase in the standard deviation of prior distribution of R_ij_ due to a unit increase in the mean of the cue q_ij_. This parameter controls how likely is a value of R_ij_ which is distant from q_ij_, compared to a value of R_ij_ which is close to q_ij_. η: The increase in the standard deviation of the prior distribution of R_ij_ due to a unit increase in the standard deviation of the cue (sd_ij_). This parameter captures an effect of the disparity (variance) in possible pain levels described in the cue on the precision of the prior. µ: The mean of the distribution of the predicted pain intensity prior to seeing the cue. ν: The standard deviation of the distribution of predicted pain intensity prior to seeing the cue.

|  | Experiment 1 |  | Experiment 2 |  |
| --- | --- | --- | --- | --- |
| Parameter | Mean | Standard Error | Mean | Standard Error |
| $\beta$ | 8.97 | 0.18 | 9.45 | 0.21 |
| $\rho$ | 6.331 | 0.37 | 8.20 | 0.45 |
| $\eta$ | 0.12 | 0.020 | 0.09 | 0.024 |
| $\mu$ | 46.05 | 0.57 | 45.61 | 0.42 |
| $\nu$ | 12.04 | 0.49 | 10.67 | 0.28 |

Table 1 also contains the DIC scores for Model 6, the ‘Hierarchical model’ where the model parameters were allowed to vary across individual. This model fitted significantly better than all fixed-parameter models (Models 1–5), suggesting that individual differences are relevant for understanding how expectation influences perceived pain. Model 6 allows us to dissect a number of aspects of the psychological construct of pain expectations, illustrated in Figures 2–4. Figure 2 illustrates the way in which Model 6 explains the responses of two different participants in Experiment 1 to a trial where these participants were exposed to the same cue. The charts on the top row illustrate the participants’ cue-independent priors before (i.e. either the cue or stimulus are delivered). In the middle row, the likelihood of the cue (red dashed curve) is incorporated to create a posterior (black curve) which will later serve as the ‘prior distribution’ (blue curve), *P* (*R_ij_*|*Z_ij_*). In the last row the ‘prior distribution’ (now the blue curve) is incorporated with the likelihood of the stimulus (red dashed curve) leading to the posterior distribution (black curve), *P*(*R_ij_*|*X_ij_*,*Z_ij_*). The left column of the figure describes a participant (participant 3) with parameters *ϐ* = 11.41, *ρ* = 13.05, *η* = 0.105, *µ* = 44.82, and *ν* = 10.00. The right column displays another participant (participant 4) with parameters *ϐ* = 11.85, *ρ* = 4.54, *η* = 0.106, *µ* = 51.23, and *ν* = 11.70. The left participant’s value for *ρ* is markedly higher than that of the right participant. This causes the ‘prior distribution’ in the second row to be markedly less precise. In the third row, this results in the posterior being closer to the actual intensity of the delivered stimulus. We might characterize the behaviour displayed in the left diagram in terms of the cue exerting little influence on the participant’s expectation and rating. The right participant’s value for *ρ* is markedly lower, which is reflected by the sharper ‘prior distribution’, centred around the cue mean, *q_ij_*. This causes the strong pull of the posterior towards the cue mean.

**Figure 2:**
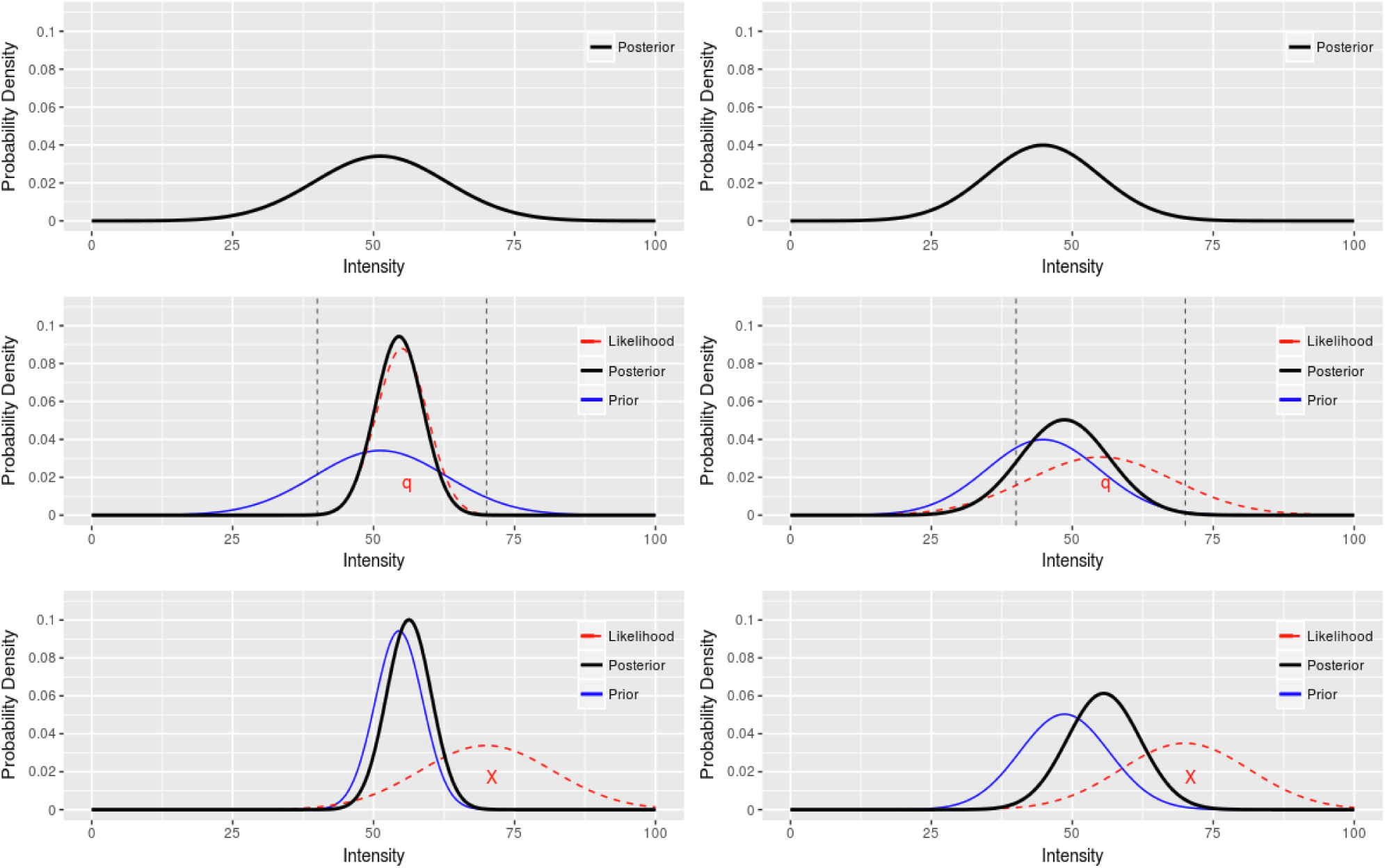
Illustration of the way the model explains the responses of two different participants in Experiment 1 after exposure to the same cue. We selected a trial in which the cue consisted of two possible pain levels (4 and 7), each with 50% probability. The horizontal axis describes pain levels on a percent scale, from 0 to 100. The two vertical bars indicate the possible levels of pain according to the cue (40 and 70). The symbol “q” marks the mean of the expected pain, based on the cue (here, 55). The symbol “X” marks the intensity X_ij_ of the delivered stimulus. Participants 3 and 4 are respectively represented in the left and right columns. The top row illustrates cue-independent prior. In the middle row, the likelihood of the cue (red dashed curve) is incorporated to create a posterior (black curve) which will later serve as the ‘prior distribution’ P(R_ij_|Z_ij_). In the last row the ‘prior distribution’ (now the blue curve) is incorporated with the likelihood of the stimulus (red dashed curve) leading to the posterior distribution (black curve), P(R_ij_| X_ij_,Z_ij_).

**Figure 3:**
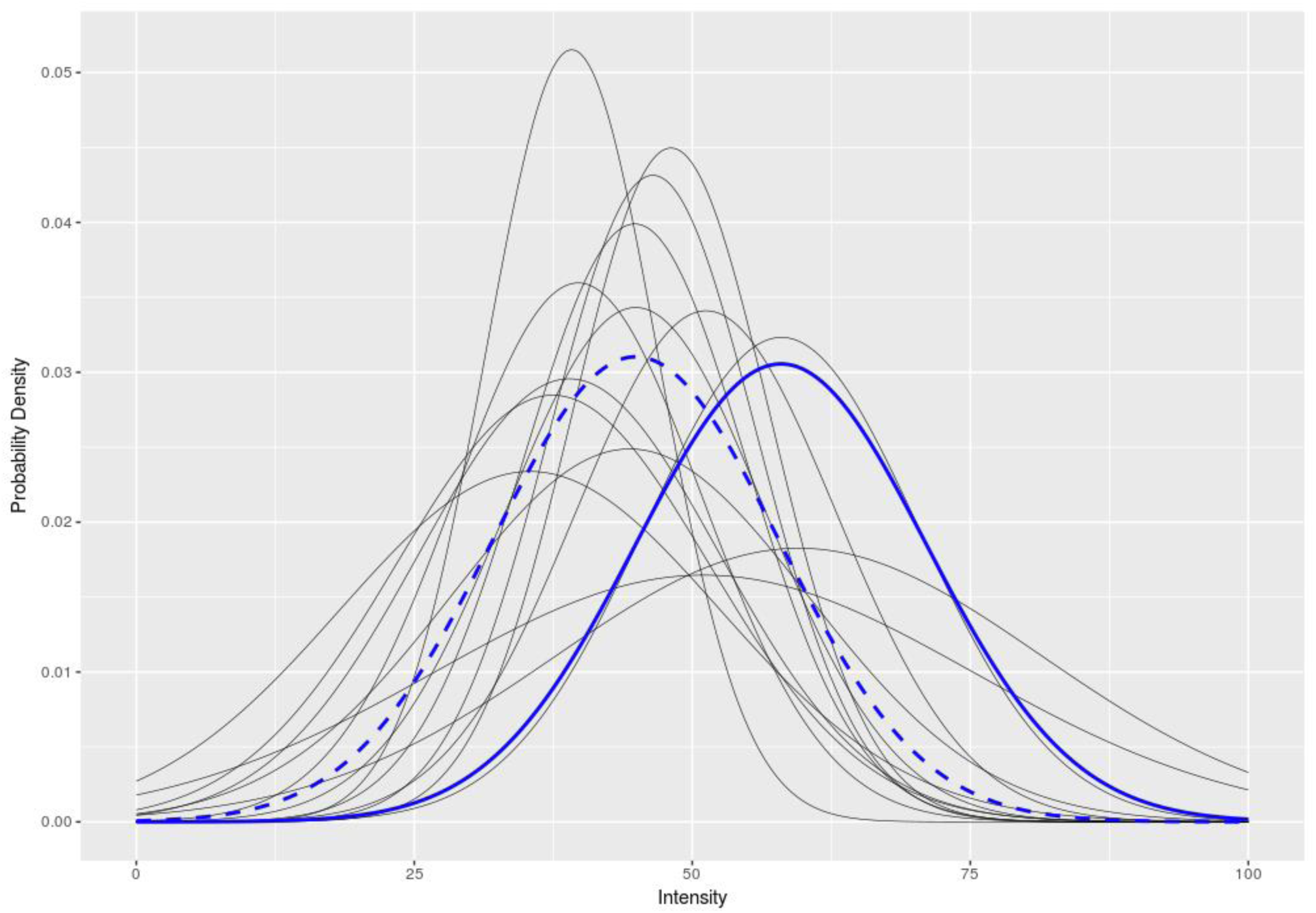
Individual-level, cue-independent priors representing trait-like bias. The graphs represent the cue-independent priors for participants in Experiment 1 under the Hierarchical model (Model 6). For example, the bold blue distribution represents a participant who is more pessimistic about pain (in terms of the mean pain that they expect) than the participant represented by the dashed blue distribution. The distributions vary in both their mean and variance with the variance determining how strongly the posterior will be biased towards the mean.

**Figure 4:**
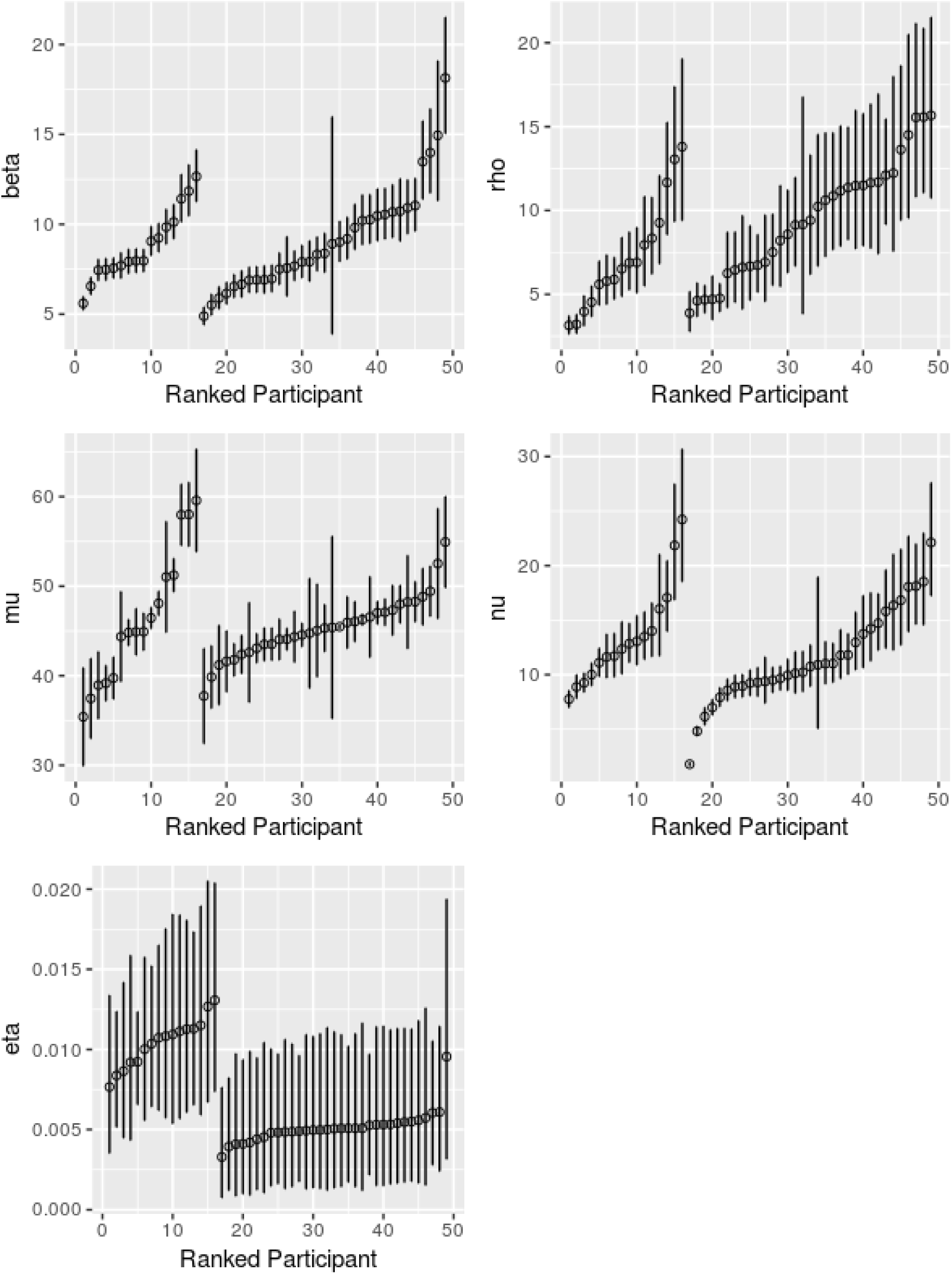
Individual-level estimates for parameters in the Hierarchical model (Model 6). There is one plot per parameter (see vertical axis label). Each plot is divided in two sections, the left section reporting the ranked estimates of the corresponding parameter for the participants in Experiment 1, and the right section reporting the ranked estimates for the participants in Experiment 2. These estimates were obtained by separately analysing the data from Experiments 1 and 2 on the basis of Model 6. Each individual in each plot is represented by a circle, whose y-coordinate corresponds to the point estimate of the parameter for that individual. Superimposed to each circle is the 95% Bayesian credible interval for the corresponding estimate. The left (respectively, right) portion of each plot refers to Experiment 1 (respectively, 2). The ordering of the individuals along the horizontal axis corresponds to their ranking by increasing value of the parameter estimate.

Figure 3 illustrates the variability in cue-independent priors among participants in Experiment 1. Here the mean and standard deviation of the curves correspond with the individual parameters µ and ν respectively. We see that the cue-independent priors differ in location and strength leading to differences in how the cue shapes the prior distribution.

Figure 4 plots the individual-specific estimates of the parameters obtained in the ‘Hierarchical model’ (Model 6) for each experiment. A clear-cut ranking of the individuals by sensitivity is evident in the plots. The estimates of parameter η appears on average to be less precise than the remaining parameters, as evident from their larger credible intervals. The individual-level estimates of the remaining parameters µ, ν, β and ρ, have narrower credible intervals, and less overlap across individuals, which suggests these parameters could provide a good basis for classifying individuals. This suggests that individuals can be accurately ranked on the basis of the latter set of parameters.

## 4. Discussion

Six models of pain perception were tested using data collected from two experiments that utilised a short, novel and deception-free experimental paradigm. A model comparison performed using data from the first experiment supported a number of hypotheses. Firstly, it was found that modelling cue-dependent priors using the mean pain intensity suggested by the cue provided a better model fit for the data than using separate modal values for each intensity represented in the cue. Secondly it was found that increased variance in the prior (and thus the generated expectation) decreased the influence which that prior exerted on the perception of pain. Thirdly, it was shown that adding parameters that describe cue-independent biases in pain perception significantly improved model fit, suggesting that such biases produce an effect on pain perception that is independent of cued pain expectations. Fourthly, as the best fitting ‘Full model’ corresponded most closely to the predictive coding framework, the results of this study lend support for the predictive coding conceptualisation of pain perception. Finally, this winning model produced a significantly better fit of the data when its parameters were allowed to vary at an individual level, suggesting that there are substantial individual differences in sensitivity to aspects of pain expectations. The results also show that this individual-level model can be used to distinguish individuals based on the characteristics of their pain expectations. Importantly, the above model comparison results were replicated when the analysis was repeated using data from a second experiment, a conceptual replication of the first which involved an independent sample.

The comparison between the fits achieved by the models presented in this paper allowed us to assess the viability of the predictive coding framework. The fact that Models 2 and 3 outperformed Model 1 replicated the typical finding the expectation can alter pain perception (e.g. Atlas & Wager, 2012; Bingel et al., 2011; Colloca, & Benedetti, 2006; Peerdeman et al., 2016; Tracey, 2010). More interestingly, the fact that Model 4 outperformed Model 3 confirms that the uncertainty around the expected pain value modulates its influence on pain perception (Brown, Seymour, Boyle, El-Deredy, & Jones, 2008b; Colloca, Petrovic, Wager, Ingvar, & Benedetti, 2010). Given that the model parameter which captures the effect that variance within the cue information has on the prior (η) was found to have a positive value, it can be concluded that greater uncertainty in the pain expectation decreases its effect on perception. This finding aligns with predictive coding, but is in contrast to previous work which has suggested that greater expectation uncertainty may increase the influence of expectation (Watkinson et al., 2013) or may increase the level of perceived pain independent of the expectation (Yoshida et al., 2013). As there were many discrepancies in the methodology used between these three studies, the reason for these divergent findings will need to be investigated in future research. It is possible that the different types of pain stimuli used (pressure by Watkinson et al., temperature by Yoshida et al.) may generate different results to those achieved via the electrical pain stimulation used in the current study. Indeed, although expectations of electric pain and laser pain influence ERP markers of pain delivery similarly (Hird et al., 2018), their effect on neural markers of pain *anticipation* differs (Babiloni et al., 2007; Hird et al., 2018). Alternatively, as Yoshida et al. used a cue which involved seeing ratings in the form of a number of ‘ticks’ on a visual-analogue scale, such that greater uncertainty was operationalised in ticks that were more spread out, there is a possibility that participants may have misinterpreted the meaning of this display, perhaps thinking that a larger spread implied greater pain intensity. It is worth noting that of all the parameters in the ‘Full model’, the estimates of parameter η appeared to be less precise than the other parameters when calculated at an individual level (Figure 4.). The values of η also differed noticeable between the two experiments. Therefore, further research is needed to understand exactly how expectation uncertainty maps onto pain experience. Nevertheless, our results corroborate predictive coding accounts of pain perception, in accordance with previous claims in favour of that framework in the same and in other modalities (Clark et al., 2008; Krol & El-Deredy, 2011; Krol & El-Deredy, 2015).

Another finding of interest in relation to the cognitive aspects of pain expectation was that Model 5 outperformed Model 4, a result which suggests that cue-independent (i.e. trait-like) differences in pain expectation have a significant effect on pain perception over and above the effect of cue-dependent pain expectations. The way the effect of cue-independent expectations was modelled, as shaping the prior distribution that was informed by the cue, was inspired by work on cognitive expectations where optimism bias modulates participants’ ability to learn from the information they are given (Garrett & Sharot, 2017). Experimental studies of the pain perception process have often ignored the influence of expectations that exist outside of those generated by the experimental paradigm, which in clinical settings are known to contribute greatly to pain experience. In the case of chronic pain patients such expectations are likely to contribute more to perceived pain than more cue-dependent expectations. Because pessimistic expectations about procedure pain could prevent patients from taking up critical preventative treatments, such as colonoscopy (Trevisani, Zelante, & Sartori, 2014) future work on pain expectation therefore needs to consider the influence that individual differences in cue-independent pain expectations have on pain perception.

While much previous research has supported the predictive coding account of pain perception (e.g. Brown et al., 2008a; Brown et al., 2008b; Brown, El-Deredy, & Jones, 2014; Clark, 2013) most of this previous research has been carried out at a population level, without paying attention to individual-level differences in the effects of expectation. To counter this shortcoming, a version of ‘Full model’ was created which allowed the model parameters to vary from one individual to another. This individual-variant, ‘hierarchical’ model was found to fit the experimental data significantly better than its fixed-parameter equivalent, suggesting that the model and its parameters were able to characterise each individual participant in terms of their sensitivity to pain expectations. This finding further suggests that these parameters capture important aspects of each individual’s response to pain. For example parameters *ρ*, ŋ and *ϐ* quantify the individual’s sensitivity to aspects of the cue (expectation precision, its variance, and the precision of their representation of the pain stimulation). Individuals with a low value of *ρ* and a high value of *ϐ* may be more sensitive to the specific information they receive about upcoming pain, so they may be more inclined to dismiss their own physical sensations in favour of expectations generated by the cue. There are hints that patients with medically-unexplained pain complaints may belong to this group (Van den Bergh, Witthöft, Petersen, & Brown, 2017). In a clinical setting, such individuals might benefit most from psychological pain therapies that concentrate on information processing biases relating to pain, as well as from training methods to increase their somatosensory sensitivity (Huque, Poliakoff, & Brown, 2017). In contrast, parameters *µ, and ν*, capture trait-like differences in pain expectation, which are independent of contextual information (e.g. the cue information). The value of parameter *µ* specifies the mean pain expected before the effect of any contextual information is considered (i.e. the cue in the experimental paradigm used here) while parameter *ν* allows us to set apart individuals with exceptionally low or high confidence in this cue-independent expectation. These cue-independent parameters can be thought of as analogous to an optimism/pessimism trait relating to future pain. In a clinical setting, individuals with a high value of *µ* and a low value of *ν* may be more likely to benefit from psychological pain therapies that concentrate on general beliefs about pain.

The individual differences that the methodology presented in this paper quantifies go beyond what is available through existing self-report instruments, such as the well-established pain catastrophizing scale (Sullivan, Bishop, & Pivik, 1995). In chronic pain, high PCS scores are associated with reports of more severe pain and reduced efficacy of physical and psychological therapies (Keefe et al., 1989). PCS and other self-report measures offer a fast, reasonably accurate, but ultimately gross and subjective, description of pain expectation and experience. They cannot explain what aspects of expectations drive changes in pain experience. Specifically, the same score on a pain scale, given by two patients, could arise through very different psychological processes. Arguably, the method to characterise individuals’ sensitivity to expectation described in the current paper goes further, to reveal the underlying cognitive mechanism which drive the impact of expectations on pain experience. It should be acknowledged that participants in the current study were not asked to report what (if any) cognitive strategies they applied during the experimental task. It is therefore not possible to assess how the underlying cognitive mechanism that is suggested by the model compares with participant’s conscious experience of engaging in the task.

In this study participants were asked to rate the intensity of the pain stimulus, rather than its unpleasantness, so as not to explicitly draw their attention to pain affect. Intensity, a sensory-discriminative property of pain, is processed in the lateral pain system, comprising the lateral thalamus and its projections to the primary and secondary somatosensory and inferior parietal cortices. In contrast, unpleasantness, an affective-motivational property, is processed in the medial pain system, comprising the medial thalamus, prefrontal cortex, insula and mid and anterior cingulate cortices (Bowsher 1957; Rainville et al 1997; Bentley et al, 2004; Kulkarni et al 2005, 2007). It has been shown that those two systems could be activated differentially by drawing attention to either the sensory or affective properties of the noxious stimulus; but that they interact to create the pain experience. Specifically, top-down anticipatory effects of expectations in a network comprising the affective system influence the appraisal of the sensory properties of pain (Brown et al, 2014). This is highlighted by the performance of Model 5, where trait-like differences in pain expectations shape the processing of the probabilistic cue, and subsequently the evaluation of the pain stimulus. The proposed model provides a means of accessing the effect of this network behaviourally, at an individual level.

The ability of the methodology described in this paper to characterise the effect of pain expectations in individuals could provide useful information in assigning treatment plans if it were applied in a clinical setting. As patients are likely to be differentially sensitive to expectations (Brown, El-Deredy, & Jones, 2014) a method, such as the one presented in this paper, which can potentially identify which patients are most sensitive to pain expectations, and which facets of pain expectations they are sensitive to, could help inform clinical decisions. For example, pain expectations are a key target in non-pharmacological therapies for pain, such as Cognitive Behavioural Therapy (CBT). CBT is effective in reducing chronic pain and pain-related disability partly because it changes patients’ pain expectations (Bingel et al., 2011). Unfortunately, CBT is not as effective as we would wish: more than half of patients with chronic pain do not benefit from CBT (Peerdeman et al., 2016; Smith et al., 2016; de Williams, Eccleston, & Morley, 2012). Put simply, if a patient can be identified as being biased to report stronger pain they may be a better candidate for CBT than a patient who reports the delivered pain more accurately; and their specific expectation sensitivity profile can further improve the specific therapeutic intervention. It is hoped that the method described in this paper may ultimately prove useful in selecting suitable candidates for pain management treatment, adding quantitative information that is less affected by introspective limitations to existing self-report questionnaires that assess pain expectations. This could facilitate the personalisation of treatment selection, an approach that is particularly important in the current climate when health services are stretched, and access to effective treatment is limited (Enck et al., 2013). In addition to providing input to clinical decisions, the methodology presented here could also be used in clinical trial research to attempt to control for placebo effects when selecting control and experimental groups. To further the goal of providing a clinical tool using our approach, the current research needs to be repeated in a variety of clinical pain populations. Such work would be required in order to establish the clinical relevance of the individual parameters of pain expectations that have been established in this study.

Prior to clinical validation, it would be beneficial for the model and associated methodology to be tested further on non-clinical samples. The samples used in the current experiments were modest in size and were derived from a relatively homogenous population (undergraduates). A large-scale validation and replication of the model and associated methodology with a more varied non-clinical population would therefore be welcome. In addition to simply validating the model, such work could also be used to answer empirical questions relating to the model variables. For example, it would be of interest to understand whether the model variables are stable over time within the same individual, and how the model variables vary within the same individual depending on the type of pain stimulation used (e.g. laser vs electrical) and the anatomical location of the stimulation. Investigations of other non-clinical determinants of pain experience (e.g. cultural differences, differences due to prior exposure to pain) could also be performed. In addition, although trial order was randomised for each participant to prevent systematic influences due to across trial learning effects, there is scope for future exploration of learning patterns. It would be interesting, for example, to examine whether the participant’s experience up to and including trial *t - 1* influences the outcome on trial *t*. Such work would inform any clinical application of the model, while also providing more general insight into human pain perception.

One limitation of our design is that the pain intensity participants received was, on average, not very different from the pain intensity that they had expected. This may be important, because key aspects of predictive coding resemble the assimilative part of assimilation-contrast theory (Hoyland, Harvey, & Sherif, 1957), which predicts that the distance between prior and the physical stimulus is crucial. Both theories agree that when the distance is small, namely, when the input is not too discrepant from our existing anchor, we will assimilate it and our experience will reflect a ‘compromise’ between the reality and our expectations. However, when the reality is very different to our expectations assimilation-contrast theory predicts that we will experience the real, discrepant stimulus as even more discrepant compared to our expectations. Our data does not currently allow us to decide whether pain experiences show such contrast effects. An additional limitation of our design is that we are currently not able to distinguish between participants who rated what they ‘really felt’ and participants who rated their pain as closer to the values of the cue because of demand characteristics. The possible influence of demand characteristics is prevalent in studies of pain experience across paradigms, including placebo designs and cue-based designs, so this caveat is not unique to our design. Future experiments using our experimental paradigm could explore this issue further, for example by manipulating the working memory load of participants, which reduce goal-directed behaviour (Otto, Gershman, Markman, & Daw, 2013). We should also emphasise that our selection of an “optimal” model has been guided by statistical criteria, a necessary requirement to achieve useful prediction of population behaviour. However, the extent to which the model reflects cognitive processes, as well as particular theories such as the predictive coding framework, should be tested further through experimental manipulations of the relevant cognitive and computational processes.

In summary, model comparison and validation yielded a number of important findings. The predictive coding framework was shown to provide an accurate description of pain perception. In particular the framework’s predictions relating to the effect of expectation uncertainty were supported. Trait-like pain expectations that exist independent of any cued pain information were shown to have an influence on pain perception that was separate and distinguishable from those provided by cued pain expectations. Finally the methodology presented in this paper was able to characterise an individual’s sensitivity to pain expectations, potentially providing the basis for a clinical tool that could guide practitioners who deliver individual psychological pain management interventions.

## Acknowledgements

CB was partially supported by the FP7 MIMOmics European Collaborative Project, as part of the HEALTH-2012-INNOVATION scheme and by the Medical Research Council. RH and DT were partially supported by ESRC First Grant ES/I010424/1. WED acknowledges the support of CONICYT, Chile, FONDECYT project 1161378 and Basal project FB0008. We thank A. Jones for inspiring this project, N. Begum and E. Hird for helpful suggestions, and A. Sparkes for collecting some of the data reported in this paper.

All authors declare no conflict of interest.

## Supplementary material

**Supplementary Table S1:**
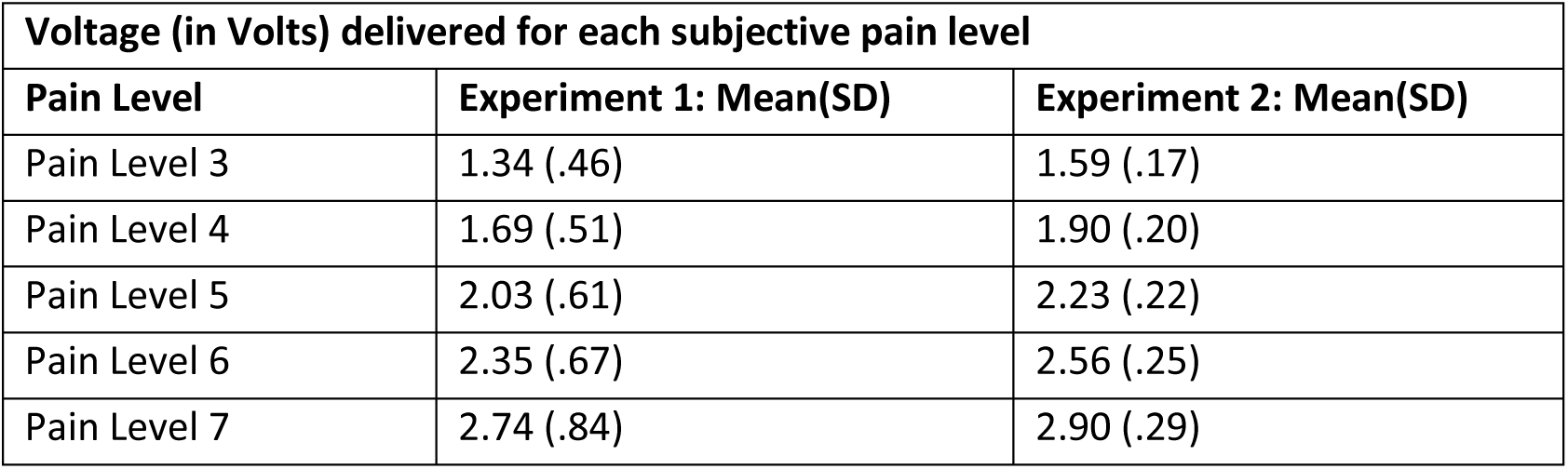
Voltage delivered for each subjective pain level in Experiments 1 and 2.

**Supplementary Table S2:** Trial types in Experiment 1.

| Possible pain levels,<br>depicted in the cue (their<br>probabilities) | Mean<br>expected<br>pain | Delivered<br>pain levels | Size of<br>Prediction<br>error at pain<br>delivery | Number of<br>repetitions<br>per block |
| --- | --- | --- | --- | --- |
| Cues with 1 option |  |  |  |  |
| 3 (100%) | 3 | 3 | 0 | 1 |
| 4 (100%) | 4 | 4 | 0 | 1 |
| 5 (100%) | 5 | 5 | 0 | 1 |
| 6 (100%) | 6 | 6 | 0 | 1 |
| Cues with 2 options |  |  |  |  |
| 4 (50%) | 4.5 | 4 | -0.5 | 1 |
| 5 (50%) |  | 5 | +0.5 | 1 |
| 3 (25%) | 4.5 | 3 | -1.5 | 1 |
| 5 (75%) |  | 5 | +0.5 | 3 |
| 4 (75%) | 4.5 | 4 | -0.5 | 3 |
| 6 (25%) |  | 6 | +1.5 | 1 |
| 5 (50%) | 5.5 | 5 | -0.5 | 1 |
| 6 (50%) |  | 6 | +0.5 | 1 |
| 4 (25%) | 5.5 | 4 | -1.5 | 1 |
| 6 (75%) |  | 6 | +0.5 | 3 |
| 5 (75%) | 5.5 | 5 | -0.5 | 3 |
| 7 (25%) |  | 7 | +1.5 | 1 |
| Cues with 4 options |  |  |  |  |
| 3 (25%) | 4.5 | 3 | -1.5 | 1 |
| 4 (25%) |  | 4 | -0.5 | 1 |
| 5 (25%) |  | 5 | +0.5 | 1 |
| 6 (25%) |  | 6 | +1.5 | 1 |
| 4 (25%) | 5.5 | 4 | -1.5 | 1 |
| 5 (25%) |  | 5 | -0.5 | 1 |
| 6 (25%) |  | 6 | +0.5 | 1 |
| 7 (25%) |  | 7 | +1.5 | 1 |

*Note.* 16 trial types correspond to a 2 (Expected pain level: 4.5, 5.5) x 2 (Discrepancy of expectations from delivered pain: 0.5,1.5) x 2 (Direction of discrepancy: positive, negative) x 2 (The number of possible pain levels participants could expect: 2, 4). In order to implement this design, 4 additional trial types were included; those are presented in light shaded boxes. For the purpose of a manipulation check, 4 further trial types with 100% probability of receiving a particular pain level were also included, presented in dark shaded boxes.

**Supplementary Table S3.**
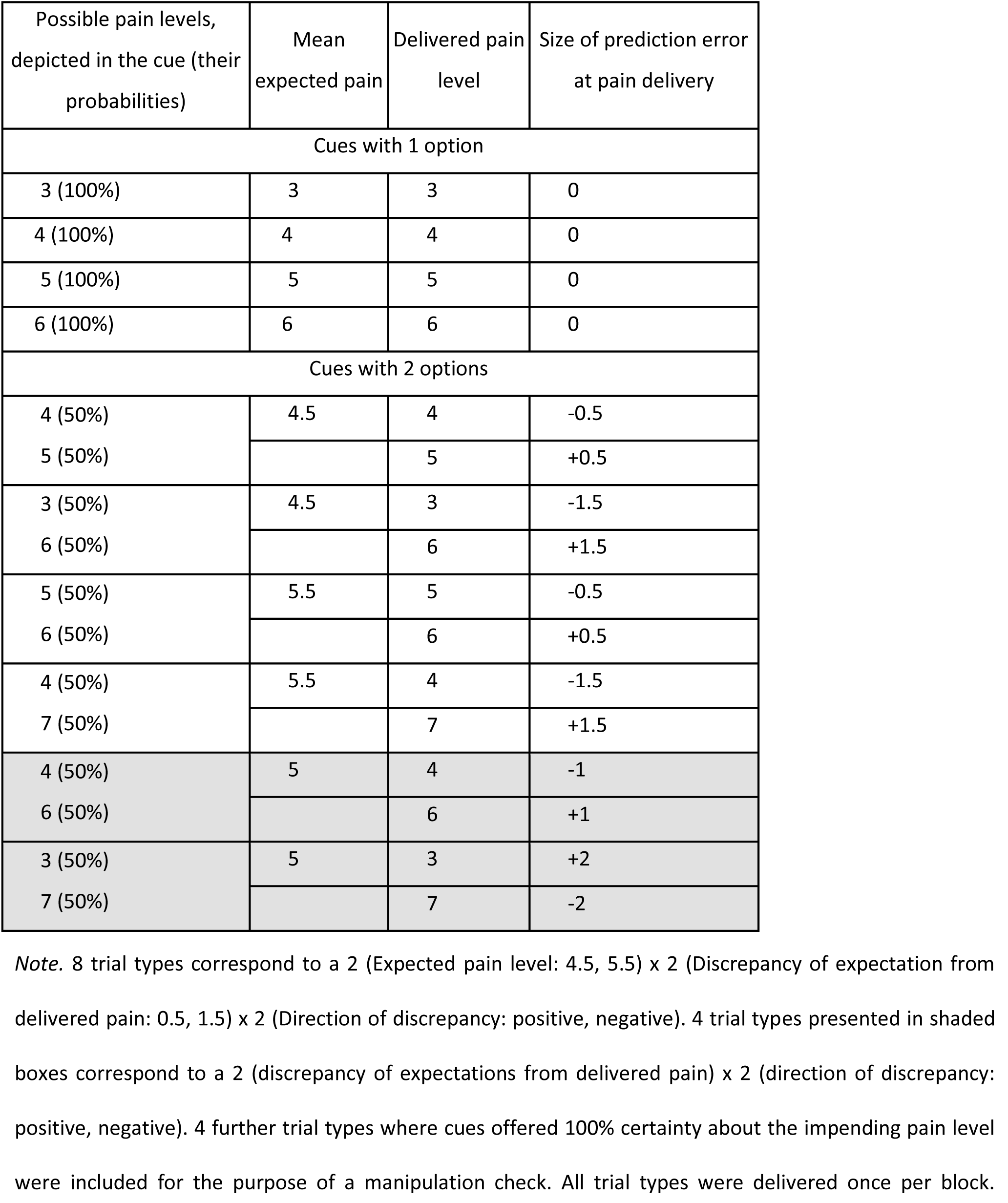
Trial types in Experiment 2.

| Possible pain levels,<br>depicted in the cue (their<br>probabilities) | Mean<br>expected pain | Delivered pain<br>level | Size of prediction error<br>at pain delivery |
| --- | --- | --- | --- |
| Cues with 1 option |  |  |  |
| 3 (100%) | 3 | 3 | 0 |
| 4 (100%) | 4 | 4 | 0 |
| 5 (100%) | 5 | 5 | 0 |
| 6 (100%) | 6 | 6 | 0 |
| Cues with 2 options |  |  |  |
| 4 (50%) | 4.5 | 4 | -0.5 |
| 5 (50%) |  | 5 | +0.5 |
| 3 (50%) | 4.5 | 3 | -1.5 |
| 6 (50%) |  | 6 | +1.5 |
| 5 (50%) | 5.5 | 5 | -0.5 |
| 6 (50%) |  | 6 | +0.5 |
| 4 (50%) | 5.5 | 4 | -1.5 |
| 7 (50%) |  | 7 | +1.5 |
| 4 (50%) | 5 | 4 | -1 |
| 6 (50%) |  | 6 | +1 |
| 3 (50%) | 5 | 3 | +2 |
| 7 (50%) |  | 7 | -2 |

*Note.* 8 trial types correspond to a 2 (Expected pain level: 4.5, 5.5) x 2 (Discrepancy of expectation from delivered pain: 0.5, 1.5) x 2 (Direction of discrepancy: positive, negative). 4 trial types presented in shaded boxes correspond to a 2 (discrepancy of expectations from delivered pain) x 2 (direction of discrepancy: positive, negative). 4 further trial types where cues offered 100% certainty about the impending pain level were included for the purpose of a manipulation check. All trial types were delivered once per block.

**Supplementary Table S4:** Rated pain experience for cues with 100% certainty in Experiments 1 and 2.

| Descriptive statistics for ratings of 100% chance trials at each pain level |  |  |
| --- | --- | --- |
| Pain level | Experiment 1: Mean (SD) | Experiment 2: Mean (SD) |
| Pain level 3 | 24.30 (8.47) | 26.43 (15.92) |
| Pain level 4 | 36.96 (10.37) | 35.25 (15.16) |
| Pain level 5 | 49.96 (13.47) | 44.59 (14.80) |
| Pain level 6 | 60.99 (16.76) | 52.72 (15.96) |

**Note: Mean (SD) of ratings on a scale of 1-100**

## Appendix

### Derivation 1: Derivation of the ‘prior distribution’

Here we derive *P*(*R_ij_*|*Z_ij_*) as resulting from a Bayesian updating process and motivate its use as the ‘prior distribution’. Following from Bayes’ theorem, the ‘prior distribution’ is proportional to the product of a cue-independent prior and a likelihood function of the cue:

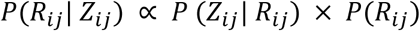

We choose the cue-independent prior to take the form:

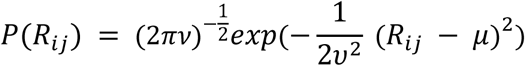

This prior is independent of the cue, and represents a stable, trait-like bias to expect pain of a certain intensity. Higher values of parameter *μ* (the mean of the cue-independent prior) reflect the expectation of higher levels of pain, independent of any cue information. Higher values of parameter *υ* (the standard deviation of the cue-independent prior) reflect greater uncertainty (and therefore lesser influence) of this cue-independent prior. Participants with high *μ* and low *υ* can be thought of as pessimistic about pain.

We choose the likelihood function of the cue to take the form:

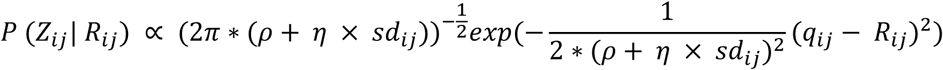

This likelihood function incorporates two parameters which will eventually modulate the effect of the cue information *Z_ij_* on the likelihood of the rating *R_ij_*. The parameter *ρ* controls the variance of the likelihood function generated by the cue. In effect it represents the precision of this prior. In contracts *η* dictates the influence the cue’s variance has on the sharpness of the likelihood function. Together these parameters determine the extent to which the cue information modulates the prior distribution, and thus the subsequent experience of pain. The influence of the cue increases with lower values of *ρ* and *η*.

Plugging these functions back into the original equation and now conditioning on *Z_ij_* we can express the prior distribution as:

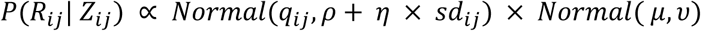

### Derivation 2: Incorporation of the Likelihood of the Stimulation

The posterior is given, up to a proportionality constant, by the product of the prior and the likelihood of the stimulation:

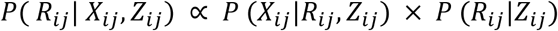

We take the likelihood of the stimulation to factor into two components, *P* (*X_ij_*|*R_ij_*,*Z_ij_*) =*f*_1_ (*X_ij_*,*R_ij_*) × *f*_2_ (*X_ij_*,*Z_ij_*). Since we consider the posterior only up to a proportionality constant, we are only concerned with the *f*_1_ (*X_ij_*,*R_ij_*) component which we take to be normally distributed:

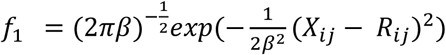
with *ϐ* > 0. Each predicted value of *R_ij_* is compared with the external sensory information, associating each such value with a measure of ‘surprise’, which our model takes to increase with an increasing distance between *R_ij_* and the actual intensity of the stimulation, *X_ij_*. The value of *ϐ* determines the extent that the perceptual mechanism will tolerate a given disparity between the predicted experience and the actual sensation, with smaller values of *ϐ* implying greater sharpness of the likelihood function around *X_ij_* and hence a reduced tolerance.

Plugging *f*_1_ back into the original equation and now conditioning on *X_ij_* we can express the posterior distribution as:

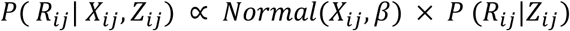

